# Rapid feedback responses are flexibly coordinated across arm muscles to support goal-directed reaching

**DOI:** 10.1101/143008

**Authors:** Jeffrey Weiler, Paul L. Gribble, J. Andrew Pruszynski

**Author notes:** Correspondence to: Jeffrey Weiler, The Brain and Mind Institute, Western University, 1151 Richmond Street, London, Ontario, Canada, N6A 3K7.

## Abstract

A transcortical pathway helps support goal-directed reaching by processing somatosensory information to produce rapid feedback responses across multiple joints and muscles. Here we tested whether such feedback responses can account for changes in arm configuration and for arbitrary visuomotor transformations – two manipulations that alter how muscles at the elbow and wrist need to be coordinated to achieve task success. Participants used a planar three degree-of-freedom exoskeleton robot to move a cursor to a target following a mechanical perturbation that flexed the elbow. In our first experiment, the cursor was mapped to the veridical position of the robot handle, but participants grasped the handle with two different hand orientations (thumb pointing upward or thumb point downward). We found that large rapid feedback responses were evoked in wrist extensor muscles when wrist extension helped move the cursor to the target (i.e., thumb upward), and in wrist flexor muscles when wrist flexion helped move the cursor to the target (i.e., thumb downward). In our second experiment, participants grasped the robot handle with their thumb pointing upward, but the cursor’s movement was either veridical, or was mirrored such that flexing the wrist moved the cursor as if the participant extended their wrist, and vice versa. After extensive practice, we found that rapid feedback responses were appropriately tuned to the wrist muscles that supported moving the cursor to the target when the cursor was mapped to the mirrored movement of the wrist, but were not tuned to the appropriate wrist muscles when the cursor was remapped to the wrist’s veridical movement.

**New and Noteworthy:** We show that rapid feedback responses were evoked in different wrist muscles depending on the arm’s orientation, and this muscle activity was appropriate to generate the wrist motion that supported a reaching action. Notably, we also show that these rapid feedback responses can be evoked in wrist muscles that are detrimental to a reaching action if a non-veridical mapping between wrist and hand motion is extensively learned.

## Introduction

Rapidly stretching a muscle of the upper-limb evokes rapid feedback responses in homonymous and/or heteronymous muscles. One such response – the long-latency stretch response – occurs approximately 50-100 ms after a muscle is stretched or mechanically perturbed. The long-latency stretch response is generated, at least in part, by a transcortical pathway that traverses regions associated with the production of voluntary movement (e.g., primary motor cortex: Cheney and Fetz 1984; Evarts and Fromm 1977; Evarts and Tanji 1976; Omrani et al, 2014; Omrani et al, 2016; Picard and Smith 1992; Pruszynski et al, 2011; Pruszynski et al, 2014, Wolpaw 1980; pre-motor and parietal cortex: Omrani et al, 2016). Likely due do this cortical processing, the long-latency stretch response can be modulated by numerous factors, including many factors that influence the production and control of voluntary movement. For example, both the long-latency stretch response and volitional movement are influenced by the intent of the action (Dimitriou et al, 2012; Colebatch et al, 1979; Crago et al, 1976; Evarts and Granit 1976; Hammond 1956; Omrani et al, 2013; Pruszynski et al, 2008), motor learning (Ahmadi-Pajouh et al, 2012; Cluff and Scott 2013), movement decision-making (Nashed et al, 2014; Selen et al, 2012; Yang et al, 2011) and environmental dynamics (Ahmadi-Pajouh et al, 2012; Kimura et al, 2006; Krutky et al, 2010). In fact, because long-latency stretch responses and voluntary actions are influenced by many overlapping factors and are generated by similar cortical circuitry, assessing how this rapid feedback response is modulated is often used as a probe of how cortical sensorimotor circuits process incoming somatosensory information to generate goal-directed actions (for reviews see Cluff et al, 2015; Pruszynski and Scott, 2012).

An important feature of the long-latency stretch response is that it can be concurrently evoked in multiple muscles, including muscles that were not stretched or mechanically perturbed. Gielen and colleagues (1988) provided a clear example of this by having participants supinate their wrist following a wrist pronation perturbation. They found that long-latency stretch responses were evoked in the biceps (a wrist supinator and elbow flexor) as well as the triceps (an elbow extensor), even though the triceps was not stretched by the perturbation. This coordination of long-latency stretch responses across multiple muscles was appropriate to stabilize the arm because biceps recruitment helps counter the perturbation but also generates unwanted elbow flexion, and this unwanted elbow flexion is countered by triceps recruitment (see also Cluff and Scott, 2013; Kurtzer et al, 2008, 2009; Pruszynski et al, 2011; Soechting and Lacquaniti, 1988). We have recently shown that long-latency stretch responses can also be evoked across multiple muscles in a coordinated way to support reaching actions. In our work, participants reached to a target following a mechanical perturbation that flexed or extended their elbow, and we placed the target in various locations where both elbow and wrist movements helped transport the hand to the desired location. We found that participants coordinated movement at both the elbow and wrist joints to complete the reaching action. More interestingly, long-latency stretch responses were not only evoked in the stretched elbow muscles but were also evoked in the wrist muscles that helped transport the hand to the target (Weiler et al, 2015; Weiler et al, 2016).

Our previous findings suggest that the rapid processing of somatosensory information accounts for how the movements of multiple joints are linked together to support reaching. Reaching to a desired location, however, can be achieved by flexibly linking the movement of several joints in numerous combinations (see Bernstein, 1967). Therefore, if somatosensory information is indeed rapidly processed to link the movements of multiple joints together to support reaching actions, this processing must ultimately account for flexible joint usage. Here we tested whether rapid somatosensory processing accounts for flexible joint usage by assessing if the long-latency stretch response evoked in wrist muscles reflects differences in how the wrist assists a reaching action.

In our first experiment we manipulated the physical orientation of the participant’s arm, which changed how wrist movement helped move a cursor to the target. For this experiment, the cursor was mapped to the veridical position of a three-degree of freedom (shoulder, elbow and wrist) exoskeleton robot handle, and participants performed reaching movements by holding the handle using two different hand orientations. In one orientation participants grasped the handle with their thumb pointing upwards (i.e., Upright Orientation), and in the other orientation participants fully pronated their forearm such that they grasped the handle with their thumb pointing downwards (i.e., Flipped Orientation). If the rapid processing of somatosensory information accounts for the arm’s orientation, then elbow flexion perturbations that displace the cursor away from the target should evoke large long-latency stretch responses in the wrist extensor muscles when the arm is in the Upright Orientation, and in the wrist flexor muscles when the arm is in the Flipped Orientation. In our second experiment we manipulated how wrist motion was mapped onto the motion of the cursor, which also changed how wrist movement helped move the cursor to the target. In this experiment, participants always held the robot handle in the Upright Orientation, but the motion of the cursor was mapped to the veridical movement of the wrist (i.e., Veridical Mapping) or to the opposite movement of the wrist (i.e., Mirror Mapping). If the rapid processing of somatosensory information accounts for the mapping between wrist and cursor movement, then elbow flexion perturbations that displace the cursor away from the target should evoke large long-latency stretch responses in the wrist extensor muscles for the Veridical Mapping, and to the wrist flexor muscles for the Mirror Mapping. Both experiments only influence how wrist movement contributes to the success of the reaching action. Therefore, if somatosensory information is rapidly processed to link the movements of multiple joints together to support goal-directed reaching, these manipulations should have limited influence on how long-latency stretch responses are evoked in elbow muscles.

## Methods

### Participants

Twenty individuals volunteered for *Experiment 1* (14 males, 6 females; mean age 21 years old) and 10 individuals volunteered for *Experiment 2* (5 males, 5 females, mean age 22). All participants reported having normal or corrected-to-normal vision and provided informed written consent prior to data collection. This study was approved by the Office of Research Ethics at Western University and was conducted in accordance with the Declaration of Helsinki.

### Apparatus

Participants grasped the handle of a three degree-of-freedom exoskeleton robot (Interactive Motion Technologies, Boston, MA). The robot allows participants to flex or extend their shoulder, elbow and/or wrist in a horizontal plane, is equipped with motors to produce flexion or extension torques at these joints and encoders to measure joint kinematics. Visual stimuli were presented downward by a 46-inch LCD monitor (60 Hz, 1,920 × 1,080 pixels, Dynex DX-46L262A12, Richfield, MN) onto a semi-silvered mirror that occluded vision of the participant’s arm. Participants were comfortably seated in a height adjustable chair and the lights in the experimental suite were extinguished for the duration of data collection.

### General Procedure

Participants began each trial by moving a cursor (turquoise circle: 1 cm diameter; see below for cursor mapping) to a red circle (i.e., the home position: 2 cm diameter) located at a position where the shoulder, elbow and wrist were at 70°, 60° and 10° of flexion (external angle coordinate system). After maintaining the cursor at this location for 1500 ms the robot applied a linearly increasing load at the elbow joint for 2000 ms that plateaued at ±3Nm (i.e., the pre-load). Participants were required to keep the cursor at the home position during this time. Note that only muscles of the elbow joint were pre-loaded by the robot. When the pre-load plateaued, the cursor was extinguished and a white target circle (10 cm diameter) was presented adjacent to the home location in one of two positions: at a location where elbow flexion would displace the hand directly into the target, or at a location where elbow flexion would displace the hand directly away the target. Participants maintained this arm position for a randomized foreperiod (1000 – 2500 ms) after which a commanded step-torque (i.e., the perturbation) of ±3Nm was applied at the elbow. Each perturbation moved the participant’s hand either into the target (IN condition) or away from the target (OUT condition) depending on the target’s location (see Figure 1B and Figure 1C). The participant’s task was to move their arm such that the cursor, if visible, would enter the target in less than 375 ms. The cursor reappeared 100 ms after the perturbation, the commanded step torque was rapidly ramped down 1000 ms after the perturbation and movement feedback was provided on each trial. If the cursor entered the target after 375 ms, or never entered the target, the target changed from white to red – otherwise the target changed from white to green.

**Fig 1.**
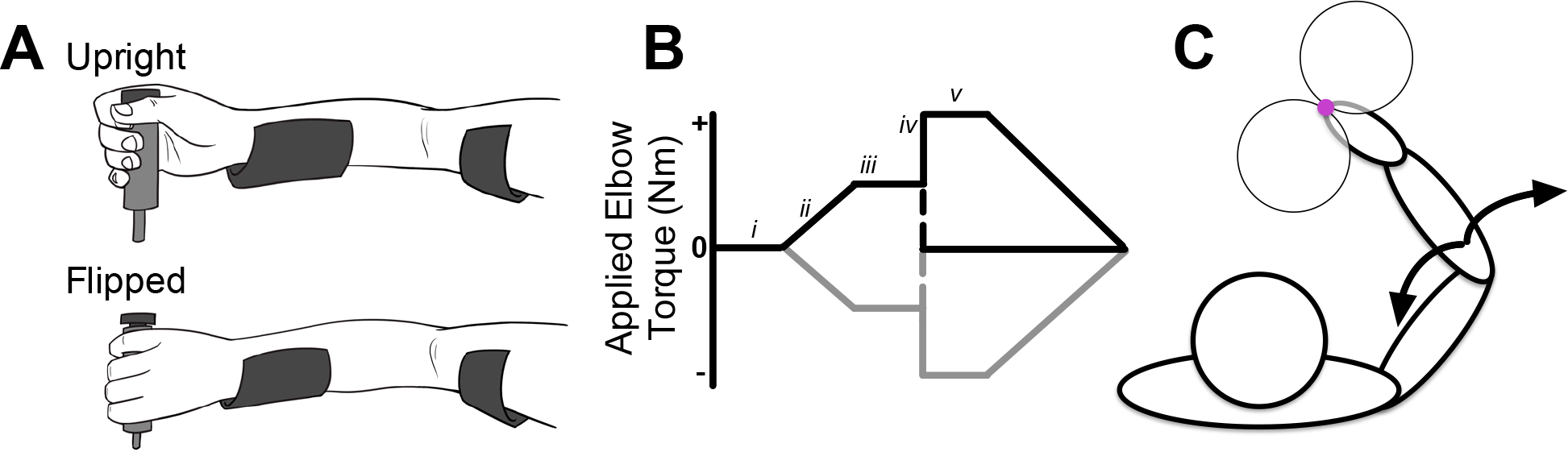
A. *Top*. Depiction of how participants grasped the exoskeleton handle for the Upright Orientation. *Bottom*. Depiction of how participants grasped the exoskeleton handle for the Flipped Orientation. B. Trial timeline *i*: Participants positioned the cursor at the home location (1500 ms). *ii*: A flexion or extension load (±3 Nm) was gradually applied at the elbow over 2,000 ms. *iii*: Visual feedback of the cursor was extinguished and white target was presented. Participants maintained this position for a randomized foreperiod (1000 – 2500 ms). *iv* A commanded step torque (±3 Nm: the perturbation) was applied at the elbow, which displaced the cursor – if it was visible – toward or away from the center of the target (IN and OUT conditions, respectively). *v* Participants moved their arm such that the cursor – if visible – moved into the target. The cursor reappeared 100 ms after the perturbation and the commanded step torque was rapidly ramped down 1000 ms after the perturbation. C. Top-down view of the experimental setup. Perturbations (i.e., commanded torques) were applied at the elbow joint (black arrows). Only one target was presented per trial.

### Mapping of the cursor

We used two different methods to map the position of the cursor relative to the robot handle. In one condition (i.e., Veridical Mapping) the cursor simply reflected the Cartesian coordinates of the robot handle. In the other condition (i.e., Mirror Mapping) the cursor reflected the Cartesian coordinates of the robot handle only when the participant’s wrist was at 10° of flexion. When participant’s wrist was not at 10° of flexion, we mapped the cursor to the Cartesian coordinates of the robot handle as if the participant flexed their wrist when they extended their wrist, and vice versa. For example, the cursor was presented as if the participant’s wrist was at 30° of extension when in fact their wrist was at 40° of flexion. We selected 10° of flexion as our reference point so that a similar arm orientation would be used to position the cursor over the home location regardless of the mapping condition (i.e., Veridical or Mirror). We used the Veridical Mapping across *Experiment 1 and 2*, whereas the Mirror Mapping was only used in *Experiment 2*.

### Experiment Specific Procedures

*Experiment 1* consisted of two blocks of trials, which differed by how participants physically grasped the handle of the robot. For one block of trials (i.e., Upright Orientation) participants naturally grasped the handle such that their forearm was in a semi-supine position (thumb pointing upwards; see top of Figure 1A). For this orientation, flexing the wrist and elbow moved the cursor in a similar direction, as did extending the wrist and elbow. For the other block of trials (i.e., Flipped Orientation) participants grasped the handle once they rotated their forearm into a fully pronated position (thumb point downwards; see bottom of Figure 1A). Notably, for this orientation, extending the elbow and flexing the wrist moves the cursor in a similar direction, as does flexing the elbow and extending the wrist. Each block consisted of 8 different trial-types (2 pre-loads: flexion, extension; 2 target locations: IN, OUT; 2 perturbations: flexion, extension). Each trial-type was repeated 30 times in a randomized order totaling 240 trials per block. The ordering of blocks was randomized across participants. Rest breaks were given approximately every 20 minutes during data collection or when requested.

*Experiment 2* took place over the course of five days and participants always grasped the robot handle with the normal Upright Orientation. For the first four days of the experiment participants practiced the same reaching task used in *Experiment 1*, but this time the movement of the cursor was mapped to the opposite movement of the wrist (i.e., Mirror Mapping). Each practice session required participants to complete 40 trials for each of the 8 trial-types (see above) in a randomized order, totaling 1280 practice trials across the four days. On the last day of the experiment participants completed two blocks of trials that differed by how the cursor was mapped with respect to the robot handle (i.e., Veridical or Mirror Mapping). Each block consisted of the 8 different trial-types, which were repeated 30 times in a randomized order totaling 240 trials per block. The ordering of blocks was randomized across participants. Rest breaks were given approximately every 20 minutes during data collection or when requested.

### Muscle Activity

Participants’ skin was cleaned with rubbing alcohol and EMG surface electrode (Delsys Bagnoli-8 system with DE-2.1 sensors, Boston, MA) contacts were coated with a conductive gel. EMG electrodes were then placed on the belly of six muscles (pectoralis major, posterior deltoid, biceps brachii long heads, triceps brachii lateral head, flexor carpi radialis, extensor carpi ulnaris) at an orientation that runs parallel to the muscle fiber. A reference electrode was placed on participants’ left clavicle. EMG signals were amplified (gain = 10^3^), and then digitally sampled at 2,000 Hz.

### Data Reduction and Analysis

Angular position of the shoulder, elbow and wrist were sampled at 500 Hz. EMG data were bandpass filtered (20 – 250 Hz, 2-pass 2^nd^-order Butterworth) and full-wave rectified. Muscle activity was normalized to their own mean activity 200 ms prior to perturbation onset when the triceps was pre-loaded by the robot (i.e., flexion pre-load). For example, the WF was normalized to its own mean activity 200 ms prior to perturbation onset when the triceps was pre-loaded. Joint kinematics and EMG were recorded from-200 ms to 400 ms relative to perturbation onset.

We analyzed kinematic and EMG data from trials in which the triceps was pre-loaded and the mechanical perturbation flexed the elbow. Trials in which the perturbation extended the elbow were excluded from analyses because these perturbations could elicit responses in the pronator teres muscle to counteract unwanted forearm supination – an action generated by biceps brachii recruitment (see Gielen et al, 1988). This is important because muscle activity from pronator teres may be interpreted as muscle activity from flexor carpi radialis, because these two muscles lie in close proximity to one another. Although not analyzed, the remaining trials were included so that participants were unable to predict what response would be required on a trial-by-trial basis.

In *Experiment 1* we estimated when wrist movement began helping transport the cursor to the target by computing time-series receiver operator characteristic (ROC) curves and fitting these curves with a segmented linear regression. For each participant, time-series ROC curves were computed from-100 – 400 ms relative to perturbation onset by using raw wrist kinematic data from IN and OUT trials within each experiment. ROC curves denote the probability an ideal observer can discriminate responses (e.g., wrist displacement) that come from two discrete categories (e.g., OUT and IN conditions), where a value of 0.5 reflects chance discrimination and value of 0 or 1 reflects perfect discrimination (Green and Swets 1966). We then fit these ROC time-series curves with a segmented linear regression to determine when these curves began to diverge from chance discrimination (~0.5). In brief, this regression technique initially finds the first of three consecutive time-series ROC samples that falls below 0.3 (or above 0.7), called x_end_, and then iteratively fits the time-series ROC curve with two linear models: model 1 is fit from x_1_ to x_i_ and is restricted to a slope of zero and model 2 is fit from x_i_ to x_end_. This process is iterated until x_i_ = x_end_ and the iteration that yields the lowest cumulative residual sum of squares between the two models is our temporal estimate of when the time-series ROC curve diverges from chance discrimination. A full description of the segmented linear regression technique, and MATLAB code for its execution, can be found in Weiler et al (2015).

Across both experiments we were primarily interested in assessing how long-latency stretch responses were modulated in wrist muscles following elbow perturbations and therefore primarily focused our analyses on the mean activity of flexor carpi ulnaris (a wrist flexor: WF) and extensor carpi radialis (a wrist extensor: WE) from 50-100 ms following the perturbation (i.e., the long-latency stretch response). WF activity from three participants in *Experiment 1* was excluded because the robot dislodged the EMG electrode during data collection. We used paired sample t-tests to compare IN and OUT condition trials for the WE and WF across both experiments. Experimental results were considered reliably different if p < 0.05. Effect sizes (Cohen’s d) are provided with all results that demonstrated reliable differences.

## Results

### Experiment 1: Changing arm orientation

The primary objective of *Experiment 1* was to test whether long-latency stretch responses evoked in wrist muscles were modulated to account for the orientation of the arm. Participants adopted one of two arm orientations (i.e., Upright or Flipped Orientation) and quickly moved the cursor into a target following a mechanical perturbation that displaced the cursor into the target (IN condition) or away from the target (OUT condition).

### Features of Elbow Behaviour

Figure 2A shows group mean elbow kinematics for the Upright and Flipped Orientation blocks. Not surprisingly, the flexion perturbation initially flexed the elbow for IN and OUT trials across the Upright and Flipped Orientation blocks. Participants then extended their elbow for OUT condition trials, which helped move the cursor towards the target. Note that the arm’s orientation had little influence on the elbow movement.

**Fig 2.**
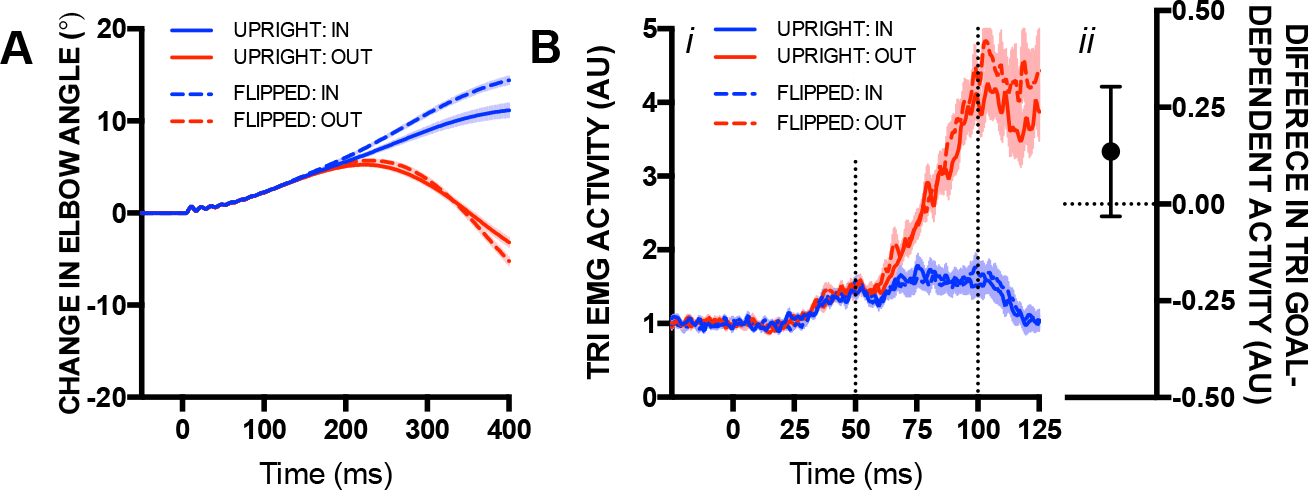
A. Mean change in elbow angle following the mechanical perturbation for trials in the Upright (solid lines) and Flipped (dashed lines) orientation. Blue and red traces reflect IN and OUT conditions, respectively. Data are aligned to perturbation onset. Shading reflects ± 1 SEM. B. *i*. Same format as A, but for mean EMG activity of the TRI. Vertical dotted lines reflect the long-latency stretch epoch. Shading reflects ± 1 SEM. *ii*. Difference in goal-dependent activity within the long-latency epoch (OUT minus IN condition) between trials completed in the Upright and Flipped orientations. Error bar denotes the 95 % confidence interval.

### Triceps long-latency stretch responses are refractory to the arm’s orientation

Figure 2B shows mean EMG activity from the triceps for the Upright and Flipped Orientations. For both arm orientations, triceps activity began to increase within the long-latency epoch for OUT condition trials relative to their IN condition counterparts. Paired sample t-tests showed that the triceps’ long-latency stretch response was larger for OUT trials compared to IN trials for both the Upright (t(19) = 8.32, p < 0.001, *d* = 1.86) and Flipped (t(19), = 9.80, p < 0.001, *d* = 2.19) Orientations. We then computed goal-dependent activity within the long-latency epoch observed in the triceps (i.e., mean activity within long-latency epoch for OUT trials minus mean activity within long-latency epoch for IN trials) for both arm orientations for each participant and compared these values with a paired-sample t-test. Results of this analysis showed no differences in triceps long-latency goal-dependent activity between the Upright and Flipped Orientations, t(19) = 1.70, p = 0.11 (Figure 2B).

### Features of Wrist Behaviour

Figure 3A displays group mean wrist kinematics for the Upright Orientation block. Note that the wrist initially moved into extension for both IN and OUT condition trials following the elbow flexion perturbation. This was expected because flexing the elbow in this arm configuration generates torque that extends the wrist due to intersegmental dynamics. The wrist then moved further into extension for OUT condition trials compared to their IN condition counterparts ~175 ms after the perturbation, which helped transport the cursor towards the target. The same basic pattern was observed when participants adopted the Flipped Orientation, but in the opposite direction. Figure 3B shows that the wrist initially moved into flexion for both IN and OUT condition trials following the elbow perturbation. This again was expected because elbow flexion in this arm orientation generates torque that flexes the wrist. The wrist moved further into flexion for OUT condition trials compared to their IN condition counterparts ~175 ms after the perturbation, which helped move the cursor towards the target. All participants displayed this general pattern of wrist motion across the two arm orientations despite not receiving explicit instructions about how to use their wrist to move the cursor towards the target. This is highlighted in Figure 3C – 3F, which shows trial-by-trial wrist motion of four exemplar participants for IN and OUT condition trials for the Upright and Flipped orientations.

**Fig 3.**
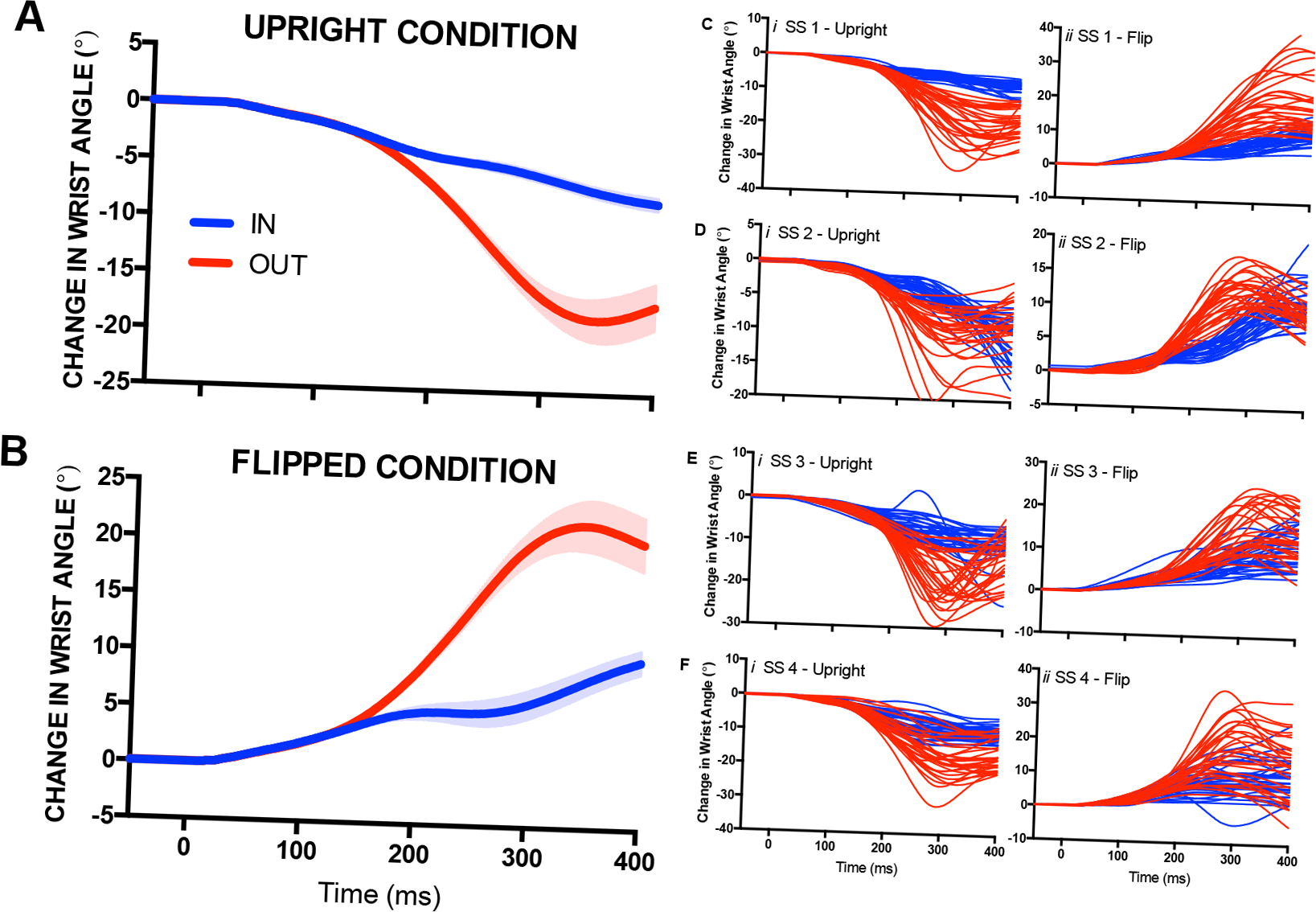
*A*. Mean change in wrist angle following the mechanical perturbation while participants adopted the Upright orientation. Blue and red traces reflect IN and OUT conditions, respectively. Data are aligned to perturbation onset. Shading reflects ± 1 SEM. *B*. Same format as *A* but for the Flipped orientation. *C-Fi*. Trial-by-trial changes in wrist angle for four exemplar participants while in the Upright orientation. Each line reflects a single trial, and blue and red traces reflect IN and OUT condition trials, respectively. Data are aligned to perturbation onset. *C-Fii*. Same format as *C-Fi*, but for the Flipped orientation.

Figure 4A shows time-series ROC curves for the Upright and Flipped Orientations from an exemplar participant each fit with our segmented linear regression technique. We computed estimates of when the time-series ROC curve diverged from chance discrimination for both the Upright and Flipped Orientations (Figure 4B). We compared these estimates with a paired sample t-test and found that the arm’s orientation did not influence when the wrist kinematics of OUT and IN condition trials began to differ, t(15) = 0.78, p = 0.49. Note that estimates of four participants were not determined because one or both of their time-series ROC curves did not fall below a value of 0.3 for three consecutive samples (see Methods).

**Fig 4.**
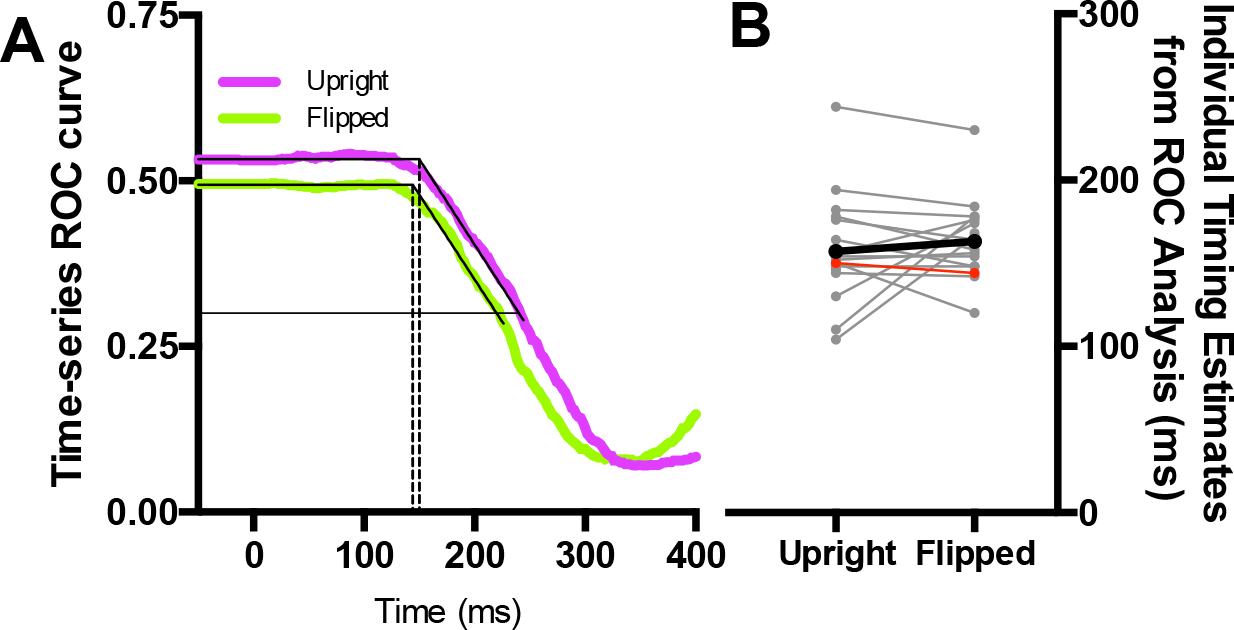
*i*. Exemplar participant’s time-series ROC curve generated for the Upright (purple trace) and Flipped (green trace) orientations. Both curves are fit with a segmented linear regression (solid black lines), which estimates the moment when the time-series ROC diverges from chance level discrimination (~0.5: vertical dashed lines). *ii*. Group estimates when the time-series ROC diverges from chance level discrimination for the Upright and Flipped Orientatoins. Thin grey lines reflect individual participants. Thin red line reflects the exemplar participant from panel *i*. Thick black line reflects the group mean.

### Flexible generation of long-latency stretch responses in wrist muscles

Figure 5 shows mean EMG activity from the WE and WF for the Upright and Flipped Orientations. For the Upright Orientation, mean EMG activity of the WE increases within the long-latency stretch epoch for the OUT compared to IN condition trials, whereas the mean EMG activity of the WF appears to be matched at all time points. For the Flipped Orientation, mean EMG activity of the WE appears to decrease within the long-latency stretch epoch for the OUT compared to IN condition trials, whereas the mean EMG activity of the WF appears to increase within the long-latency stretch epoch for the OUT compared to IN condition trials.

**Fig 5.**
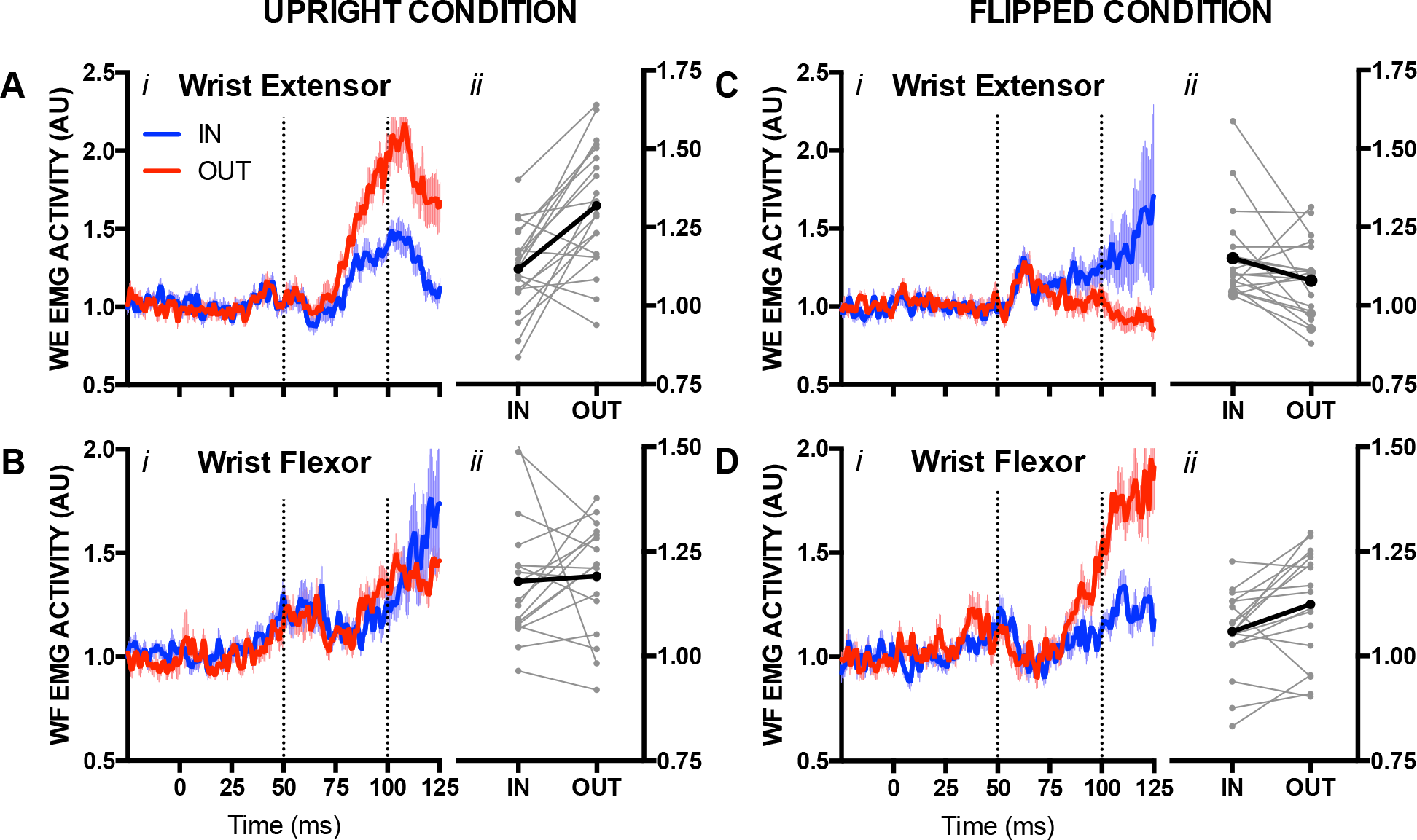
*Ai*. Mean EMG activity of the WE following the mechanical perturbation while participants adopted the Upright orientation. Blue and red traces reflect IN and OUT conditions, respectively. Data are aligned to perturbation onset. Vertical dotted lines reflect the long-latency stretch epoch. Shading reflects ± 1 SEM. *Aii*. Mean EMG activity of the WE in the long-latency stretch epoch for IN and OUT conditions while participants adopted the Upright orientation. Thin grey lines reflectindividual participants whereas thick black lines reflect the group mean. *Bi*. Same format as *Ai*, but for the WF. *Bii*. Same format as *Aii*, but for the WF. *Ci*. Same format as *Ai*, but for the Flipped orientation. *Cii*. Same format as *Aii*, but for the Flipped orientation. *Di*. Same format as *Bi*, but for the Flipped orientation. *Dii*. Same format as *Bii*, but for the Flipped orientation.

We used paired sample t-tests to compare mean EMG activity within the long-latency stretch epoch between IN and OUT condition trials from the WE and WF for both the Upright and Flipped Orientation. For the Upright Orientation, we found that WE long-latency stretch responses were larger for OUT condition trials compared to their IN condition trial counterparts, t(19) = 4.70, p < 0.001, *d* = 1.05, whereas there was no reliable difference in long-latency stretch responses between IN and OUT condition trials for the WF, t(16) = 0.29, p = 0.78. For the Flipped Orientation, we found that WE long-latency stretch responses were smaller for OUT condition trials compared to their IN condition trial counterparts, t(19) = −2.11, p = 0.041, *d = 0.47*, and that WF long-latency stretch responses were larger for OUT condition trials compared to their IN condition trial counterparts, t(16) = 2.72, p = 0.015, *d* = 0.66.

### Experiment 2: Changing visual mapping

The objective of *Experiment 2* was to test whether long-latency stretch responses evoked in wrist muscles were modulated to account for a non-veridical mapping between wrist movement and cursor movement. To test this, we mapped the movement of the cursor such that it moved as if the participant extended their wrist when if fact they flexed their wrist, and vice versa (i.e., Mirror Mapping). Thus, if an elbow flexion perturbation displaced the cursor away from the target, participants would have to flex their wrist – not extend their wrist – in order to help transport the cursor to the target’s location. Our initial pilot testing, however, demonstrated that participant’s were unable to account for this mapping within a single experimental session. We therefore trained individuals for four days to complete the reaching task with the Mirror Mapping, and then assessed long-latency stretch responses in the wrist muscles on the 5^th^ day.

### Features of Elbow Behaviour

Figure 6A shows group mean elbow kinematics for the four practice days of the Mirror Mapping condition, as well as the final day of testing, in which participants completed both the Veridical and Mirror Mapping conditions. As was observed in *Experiment 1*, the mechanical perturbation initially flexed the elbow for all trial-types across the Veridical and Mirror Mapping conditions. Participants then extended their elbow for OUT condition trials, which helped transport the cursor to the target. Note that Figure 6 shows that the pattern of elbow movement was the same across all testing sessions, and was not substantially altered by how the cursor was mapped with respect to motion of the wrist.

**Fig 6.**
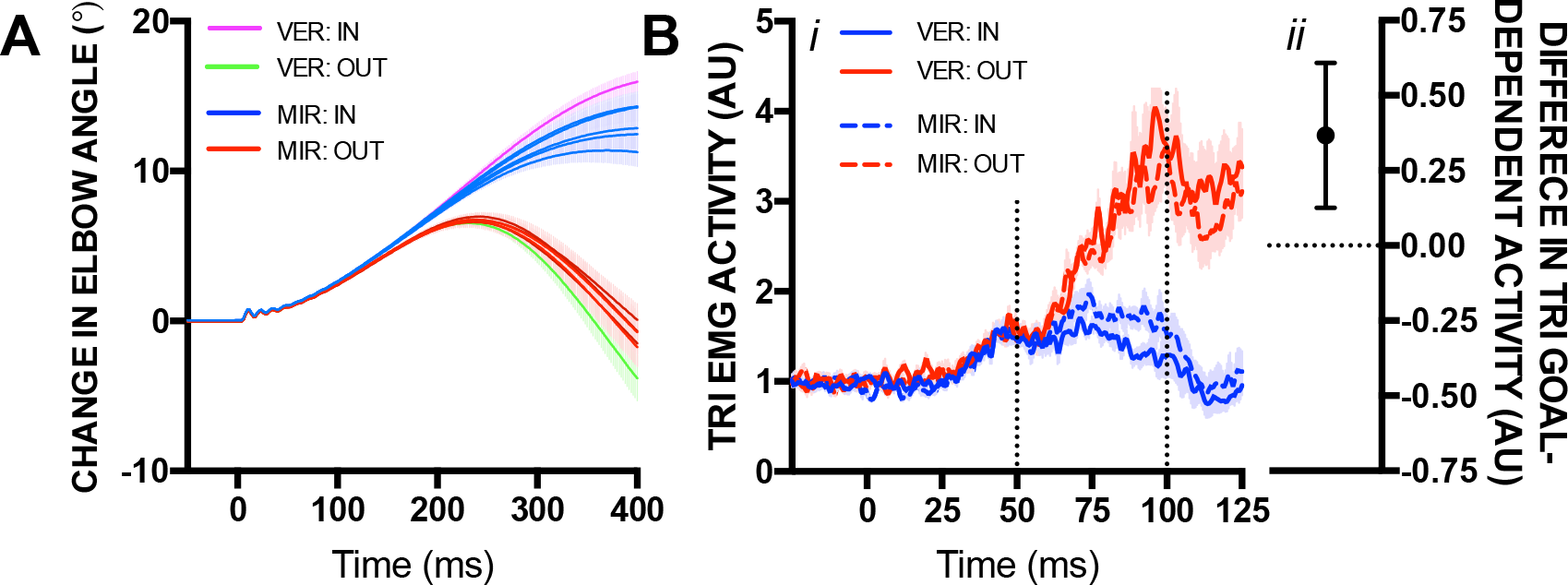
A. Mean change in elbow angle following the mechanical perturbation for trials completed across the four days of practice and the final testing session for the Mirror Mapping condition (IN: blue; OUT: red) and the Veridical Mapping condition (IN: purple; OUT: green). Data are aligned to perturbation onset. B. *i*. Mean EMG activity of the TRI on the final testing session. Solid and hashed lines reflect the Vertical Mapping and Mirror Mapping conditions, respectively, and blue and red traces reflect IN and OUT trials, respectively. Vertical dotted lines reflect the long-latency stretch epoch. Shading reflects ± 1 SEM. *ii*. Difference in goal-dependent activity within the long-latency epoch (OUT minus IN condition) between trials completed in the Veridical and Mirror Mapping conditions. Error bar denotes the 95 % confidence interval.

### Triceps long-latency stretch responses are refractory to the mapping of the cursor

Figure 6Bi shows mean EMG activity from the triceps for the Veridical and Mirror Mapping conditions. Regardless of how the cursor was mapped with respect to the wrist, triceps activity began to increase within the long-latency epoch for OUT condition trials relative to their IN condition counterparts. Paired sample t-tests showed that the triceps long-latency stretch response was larger for OUT trials compared to IN trials for both the Veridical (t(9) = 6.48, p < 0.001, *d* = 2.05) and Mirror (t(9), = 6.60, p < 0.001, *d* = 2.09) Mapping conditions. We again computed goal-dependent activity within the long-latency epoch from the triceps for trials completed in the Veridical and Mirror Mapping blocks for each participant and compared these values with a paired sample t-test. Results of this analysis showed that there was more goal-dependent activity for trials completed in the Veridical Mapping condition compared to the Mirror mapping condition, t(9) = 3.42, p = 0.007, *d* = 1.08 (Figure 6Bii).

### Features of Wrist Behaviour

Figure 7 shows three exemplar participants’ trial-by-trial wrist kinematics, as well as group mean wrist kinematics, across the four days of practice and the final testing session. Recall from *Experiment 1* that, in the Upright Orientation, participants’ wrists initially moved into extension for both IN and OUT condition trials following the elbow flexion perturbation. The wrist then further extended for OUT condition trials to help move the cursor towards the target. In *Experiment 2*, participants’ wrists also initially moved into extension for both IN an OUT condition trials following the elbow flexion perturbation. However, because the cursor’s movement was mirror mapped with respect to the wrist, participants had to flex their wrist – not extend their wrist – to help move the cursor to the target. As shown in Figure 7, participants frequently made inappropriate extension wrist movements, or delayed wrist flexion movements, for OUT condition trials during the initial training sessions. Participants progressively improved over the four days of practice and were consistently using their wrist in an appropriate fashion to move the cursor towards the target on the final testing day. This is highlighted in Figure 8A, which shows the mean movement time (MT) for participants to rotate their wrist 10 degrees in a direction that moved the cursor towards the target across the four days of practice of the Mirror Mapping condition, the final testing day of both the Mirror and Veridical Mapping conditions, as well as the Upright and Flipped Orientations from *Experiment 1*. As shown in the figure, the MT on the final testing session for the Mirror Mapping was comparable to the MTs from the other experimental conditions (i.e., Veridical Mapping, Upright and Flipped Orientation). These practice sessions also resulted in an increase in task performance. Figure 8B depicts the percentage of responses where participants received ‘successful’ feedback for OUT condition trials, which increased as a function of practice. This increase in task success was most likely attributed to how participants learned to use their wrist over the course of practice, because elbow responses remained fairly consistent over this time period (see Figure 6A).

**Fig 7.**
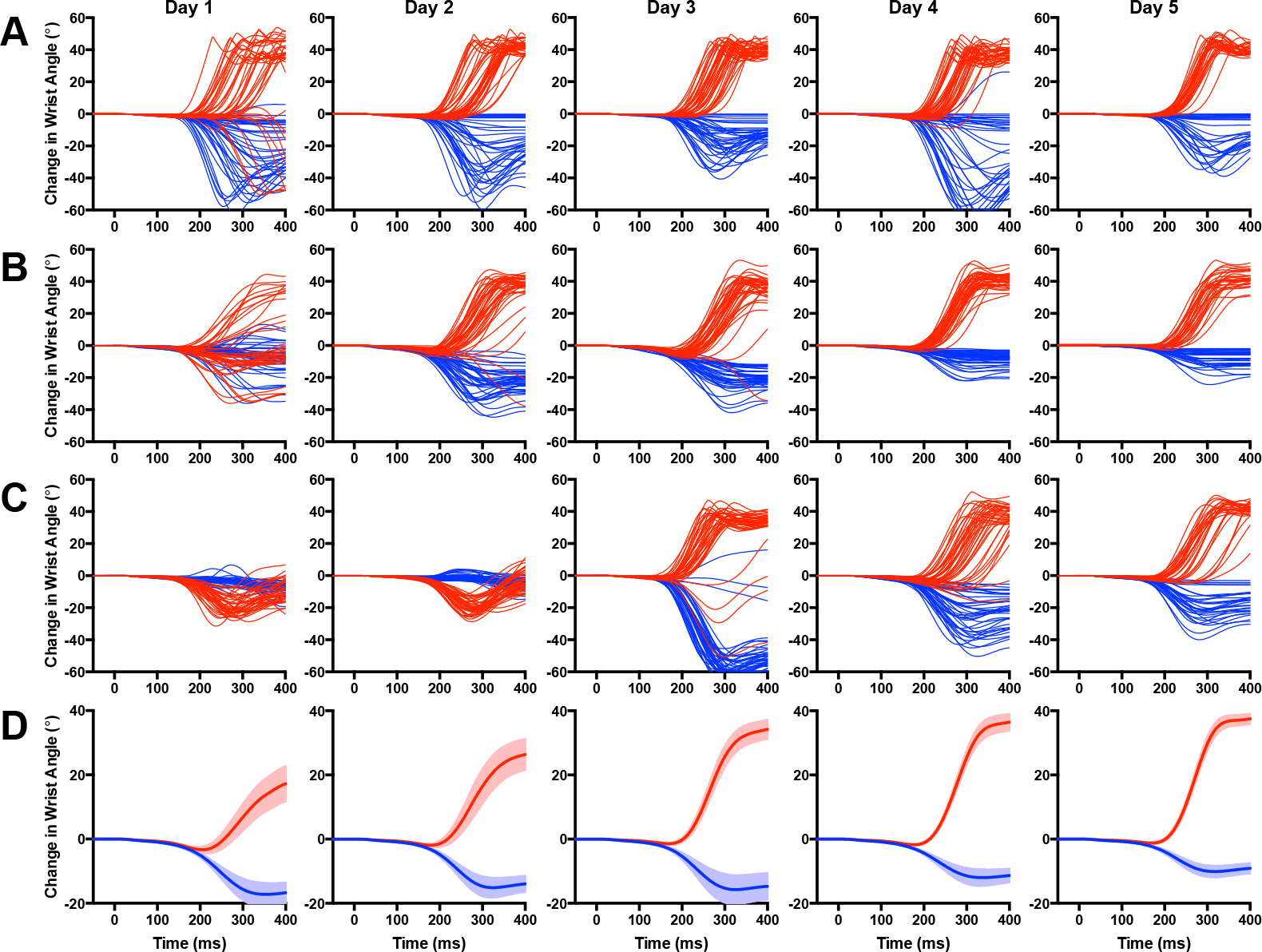
*A-C*: Trial-by-trial changes in wrist angle from three exemplar participants for the Mirror Mapping block across the four days of practice and the final training session. Each line reflects a single trial, and blue and red traces reflect IN and OUT condition trials, respectively. Data are aligned to perturbation onset. *D*: Mean change in wrist angle following the mechanical perturbation for the Mirror Mapping block across the four days of practice and the final training session. Blue and red traces reflect IN and OUT conditions, respectively. Data are aligned to perturbation onset. Shading reflects ± 1 SEM.

**Fig 8.**
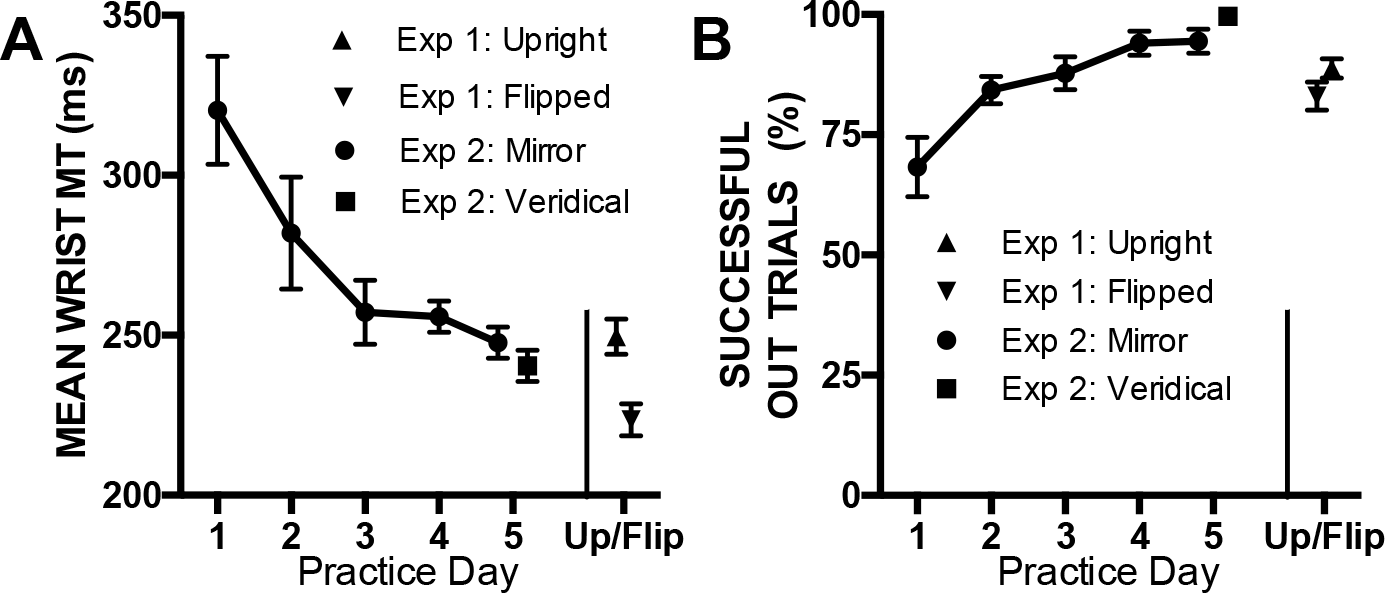
A Mean movement time required for participants to rotate their wrist 10 degrees in a direction that brings the cursor towards the target following a perturbation that displaced the cursor away from the target. Connected circles denote the required movement times over the course of the Mirror Mapping training session and final day of testing, whereas the square, triangle and diamond reflect the movement time required for the Veridical Mapping, Upright Orientation and Flipped Orientation, respectively. Error bars reflect 1 SEM. B. Same format as A, but for the proportion of OUT trials where participants successfully reached the target in less than 375 ms after perturbation onset.

### Flexible but partially erroneous generation of long-latency stretch responses in wrist muscles

Mean EMG responses of the WE and WF from the Veridical and Mirror Mapping blocks on the final testing day are shown in Figure 9. For the Mirror Mapping block, WE long-latency stretch responses did not differ between OUT and IN condition trials, t(9) = 1.00, p = 0.34, whereas WF long-latency stretch responses for OUT condition trials were reliably larger than IN condition trials, t(9) = 4.65, p = 0.001, *d* = 0.46. These results are consistent with the idea that the mechanism that generates the long-latency stretch response can account for the non-veridical mapping between wrist and cursor. For the Veridical Mapping block, WE long-latency stretch responses for OUT conditions were reliably larger than their IN condition counterparts, t(9) = 4.65, p < 0.001, *d* = 0.2. Notably, and in contrast to *Experiment 1* as well as our previous work (Weiler et al, 2015; Weiler et al, 2016), WF long-latency stretch responses were also larger for OUT condition trials compared to IN condition trials, t(9) = 2.85, p = 0.02, *d* = 0.9. This was surprising, as this pattern of WF activity is inappropriate to produce wrist extension, which is the wrist movement that would help move the cursor towards the target.

**Fig 9.**
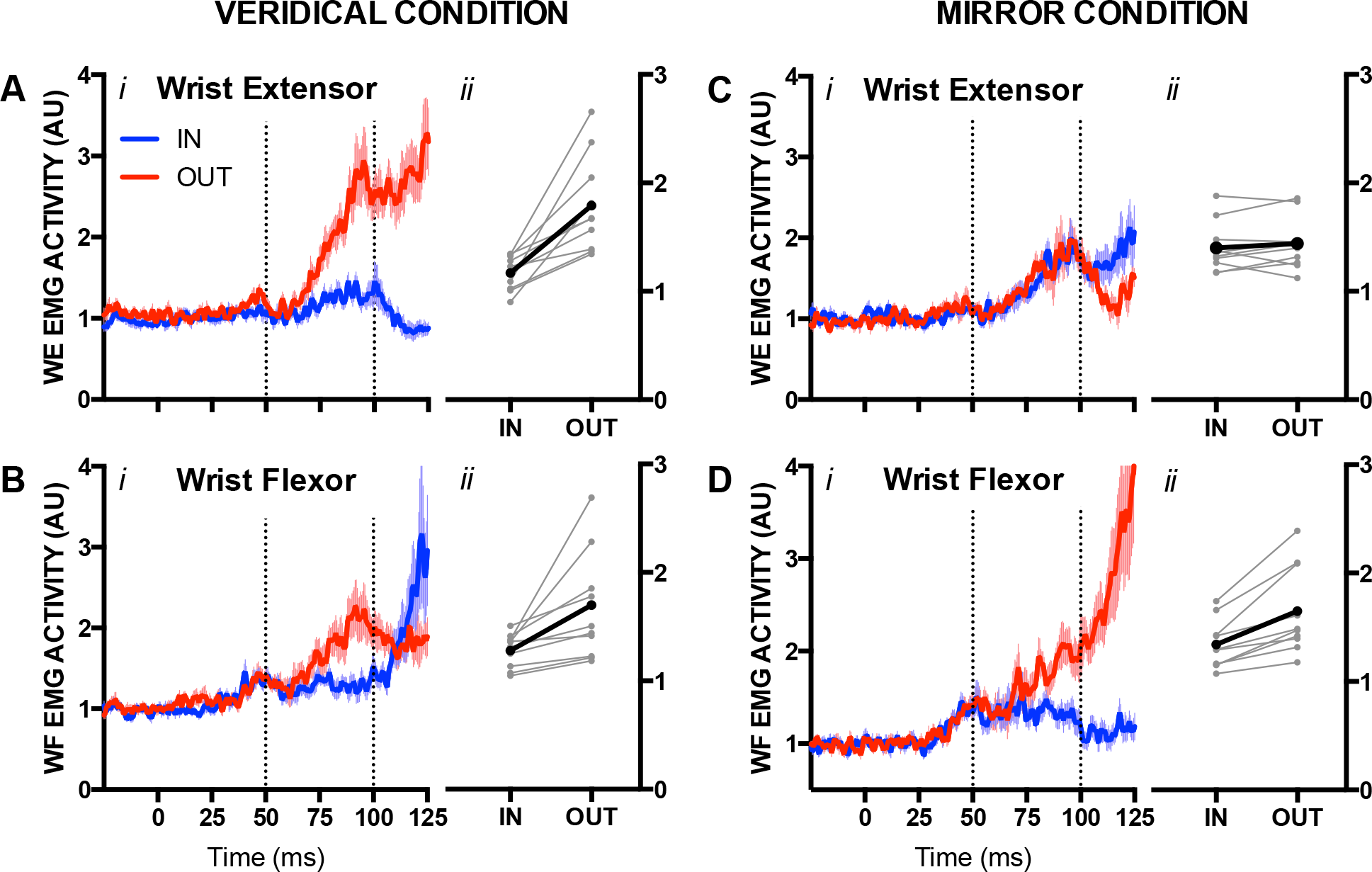
*Ai*. Mean EMG activity of the WE following the mechanical perturbation for the Veridical Mapping block. Blue and red traces reflect IN and OUT conditions, respectively. Data are aligned to perturbation onset. Vertical dotted lines reflect the long-latency stretch epoch. Shading reflects ± 1 SEM. *Aii*. Mean EMG activity of the WE in the long-latency stretch epoch for IN and OUT conditions for the Veridical Mapping block. Thin grey lines reflect individual participants whereas thick black lines reflect the group mean. *Bi*. Same format as *Ai*, but for the WF. *Bii*. Same format as *Aii*, but for the WF. *Ci*. Same format as *Ai*, but for the Mirror Mapping block. *Cii*. Same format as *Aii*, but for Mirror Mapping block. *Di*. Same format as *Bi*, but for Mirror Mapping block. *Dii.* Same format as *Bii*, but for the Mirror Mapping block.

One possible explanation for this unexpected result is that the extensive training of the Mirror Mapping block conditioned WF long-latency stretch responses to be elicited independent of the participants’ volitional movement. We tested this idea by comparing the WF long-latency stretch responses for OUT condition trials between the Veridical and Mirror Mapping blocks, as well as mean EMG activity within the voluntary epoch (i.e., 100-300 ms following perturbation onset) for OUT condition trials between the Veridical and Mirror Mapping blocks (see Figure 8). Note that the need to flex the wrist following an elbow flexion perturbation for OUT condition trials is different between the Veridical and Mirror Mapping blocks – in the Veridical Mapping block one should extend the wrist, whereas in the Mirror Mapping block one should flex the wrist. Consistent with these requirements, we found that WF muscle activity in the voluntary epoch was larger for OUT condition trials in the Mirror Mapping block compared to the Veridical Mapping block, t(9) = 6.24, p < 0.001, *d* = 1.97 (see Figure 10*ii*). In contrast, long-latency stretch responses of the WF for OUT condition trials in the Mirror Mapping block did not reliably differ from the Veridical Mapping block, t(9) = 0.19, p = 0.85 (see Figure 10*i*). Thus, long-latency stretch responses evoked in the WF within the Veridical Mapping block were similar to how these responses were evoked in the WF within the Mirror Mapping block, and did not support how participants voluntarily moved their wrist to help transport the cursor to the target.

**Fig 10.**
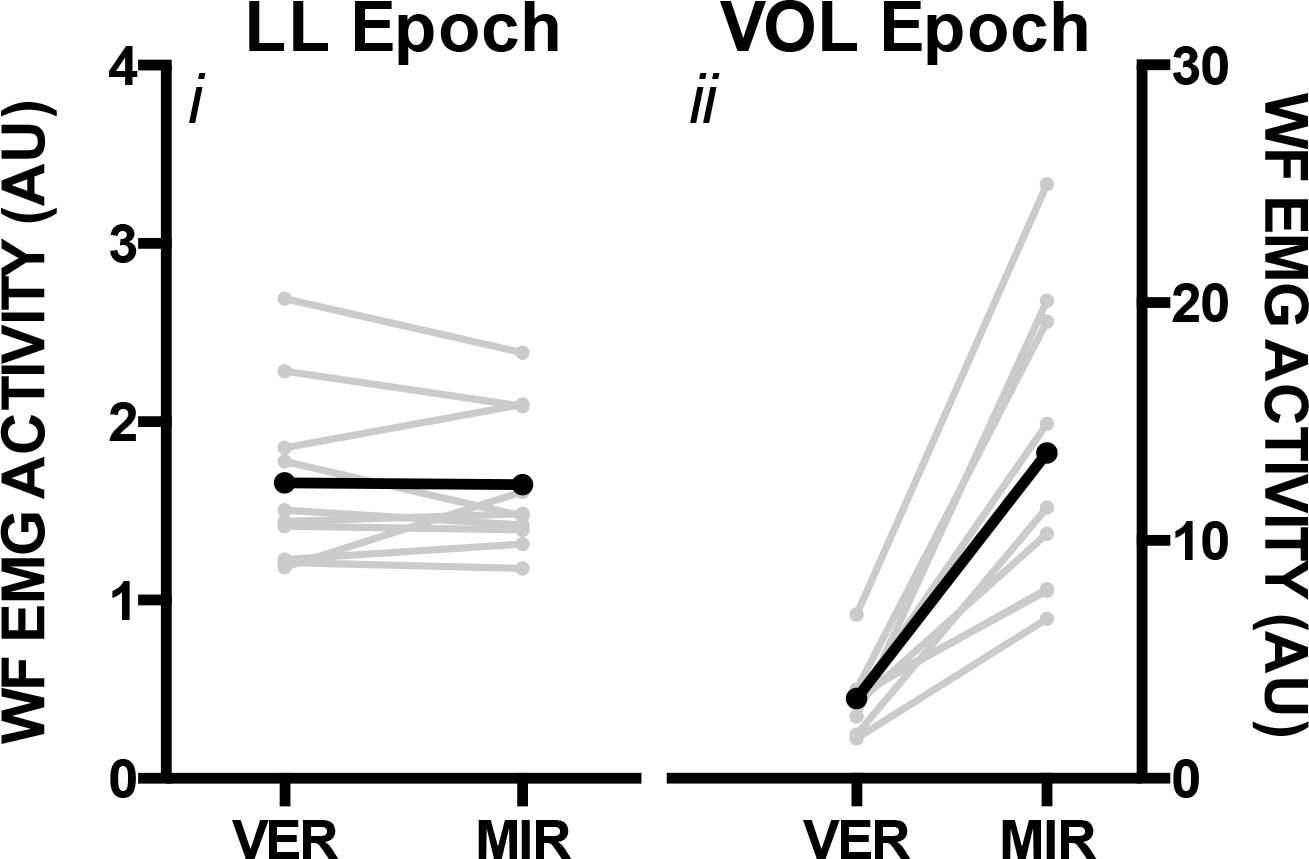
*i* Mean EMG activity of the WF in the long-latency stretch epoch for OUT condition trials between the Veridical Mapping block (VER) and Mirror Mapping block (MIR). Thin grey lines reflect individual participants whereas thick black lines reflect the group mean. *ii*. Same format as *i*, but for mean EMG activity within the voluntary epoch (i.e., 100-300 ms following perturbation onset). Note the different scales for long-latency and voluntary epochs.

## Discussion

We designed two experiments to test the flexibility with which long-latency stretch responses are generated in wrist muscles to support a simple goal-directed reaching action. In our first experiment, participants physically changed how they grasped the handle of a robotic exoskeleton, which in turn changed how the wrist moved the cursor towards the target. In our second experiment, we changed how wrist movement was mapped to the movement of the cursor. We observed three novel findings when elbow perturbations displaced the cursor away from the target. First, long-latency stretch responses were evoked in different wrist muscles depending on the arm’s orientation, and this muscle activity was appropriate to generate the wrist motion that helped transport the cursor to the target. Second, once participants learned the wrist motion required for the Mirror Mapping condition, long-latency stretch responses were evoked in a manner appropriate for moving the cursor to the target. And third, following extensive training in the Mirror Mapping condition, long-latency stretch responses were evoked in muscles that were inappropriate for bringing the cursor to the goal target when the cursor was mapped to the wrist’s veridical movement. Below, we discuss each of these points and their implications for the neural control of movement.

### Coordinating rapid feedback responses across multiple muscles for goal-directed movement

Long-latency stretch responses are generated, at least in part, by processing somatosensory information within cortical regions involved in the production of voluntary motor commands (e.g., primary motor cortex: Cheney and Fetz 1984; Evarts and Fromm 1977; Evarts and Tanji 1976; Omrani et al, 2014, Omrani et al, 2016; Picard and Smith 1992; Pruszynski et al, 2011; Pruszynski et al, 2014; pre-motor cortex and parietal cortex: Omrani et al, 2016). Therefore, assessing how these rapid feedback responses are generated across multiple muscles provides insight into how cortical sensorimotor circuits process incoming somatosensory information to flexibly coordinate purposeful motor output. The general goal of our work is to understand how somatosensory information is rapidly processed to support reaching actions and we have often approached these questions by applying mechanical perturbations to the elbow and assessing how long-latency stretch responses are evoked in multiple muscles of the upper-limb. One interesting finding we and others have extensively documented is that long-latency stretch responses are evoked in shoulder muscles to counteract torques that are generated at the shoulder joint as a result of rapid elbow motion (Crevecoeur et al, 2012; Kurtzer et al, 2008, Kurtzer et al, 2009; Maeda et al, 2017; Pruszynski et al, 2011; Soechting and Lacquaniti 1988). Such a finding indicates that the neural circuits that generate the long-latency stretch response use an internal model that accounts for biomechanical complexities (e.g., interaction torques) associated with controlling a multi-joint linkage like the arm (Kurtzer et al, 2008).

More recently, we have shown that individuals utilize elbow and wrist movement to move their hand to a goal-location following an elbow perturbation, and that long-latency stretch responses are not only evoked in the stretched elbow muscles, but also evoked in the wrist muscles that support this wrist behaviour (Weiler et al, 2015; Weiler et al, 2016). Here, we asked whether the rapid processing of somatosensory information that generates the long-latency stretch responses accounts for how movements of multiple joints can be flexibly linked together to support goal-directed reaching. In *Experiment 1* we altered the physical orientation of the participant’s arm as a direct and natural way to test this question. Participants did not alter their elbow responses but easily modified their wrist behaviour when they adopted the Flipped Orientation to help transport the cursor towards the target and our time-series ROC analysis indicated that the arm’s orientation did not influence the timing of incorporating wrist movement into the reaching action. These behavioural findings are reminiscent of Scholtz’s (2000) pistol-shooting experiment, which showed that individuals account for the initial orientation of the arm to flexibility coordinate the movement of multiple joints when executing a volitional action. What was novel about our current results was that large long-latency stretch responses were always evoked in the triceps, but flexibility evoked in the WF when wrist flexion assisted the reaching movement and in the WE when wrist extension assisted the reaching movement, and that this flexibility required no training or practice. Thus, both voluntary actions and rapid feedback responses possess the ability to flexibly coordinate the movement of multiple arm joints during reaching, possibility because they both have access to a similar – or the same – internal model that accounts for the arm’s dynamics.

There are several regions that could perform the computations of an internal model, by appropriately tuning or generating rapid feedback responses to support reaching actions. Primary motor cortex is one potential candidate. Neurons in this region respond ~20ms after mechanically perturbing a variety of upper-limb joints (shoulder: Omrani et al, 2014; Omrani et al, 2016; Pruszynski et al, 2011; elbow: Evarts and Tanji, 1976; Omrani et al, 2014; Omrani et al, 2016; Pruszynski et al, 2011: wrist: Cheney and Fetz, 1984), and then modify their firing rates ~30 ms later based on the demands of the task (Omrani et al, 2014; Omrani et al, 2016; Pruszynski et al, 2014; see also Evarts and Tanji 1976). The initial perturbation evoked activity that transitions to task or goal-dependent activity may reflect the computations of an internal model that is housed within the local circuitry of primary motor cortex. Alternatively, it is possible that the 30 ms generic-to-goal-dependent transition results from computations performed in other regions, which then modify the activity of primary motor cortex neurons. The cerebellum is another candidate region for these types of computations. Firing rates of neurons in the dentate nucleus are modulated by different movement goals ~30 ms following a mechanical perturbation (Strick 1983) and cooling this region delays the generic-to-goal-dependent transition of firing of neurons in primary motor cortex (Vilis and Hore, 1980). Thus, it is possible that the initial perturbation related activity observed in primary motor cortex is routed to the cerebellum, and the output of this cerebellar processing is rerouted back to primary motor cortex to modulate the activity of neurons that contribute to the generation of rapid feedback responses.

### Motor learning and rapid somatosensory processing

A feature of *Experiment 2* was that participants required several days of practice to learn the non-veridical relationship between the movement of wrist and motion of the cursor. Once this difficult relationship was learned, large long-latency stretch responses were evoked in the WF muscles when the perturbation displaced the cursor away from the target (i.e., OUT trials). This pattern of muscle activity was appropriate for trials in the Mirror Mapping condition because it produced the wrist flexion movement that helped transport the cursor to the target. This result indicates that long-latency stretch responses can be modulated to account for a novel visuomotor transformation, and highlights a linkage between learning novel movements and how somatosensory information is rapidly processed. A similar linkage has been previously shown for force field adaptation. For example, Cluff and Scott (2013) had participants reach to targets that required shoulder and/or elbow movement and applied a viscous force field that opposed elbow motion. On a subset of trials a mechanical perturbation rapidly extended the elbow when the participant initiated their reach. Perturbations applied when participants reached to targets evoked long-latency stretch responses in the stretched elbow muscle, but critically, the gains of these rapid feedback responses were dependent on how well the individual learned to counteract the viscous force field when reaching to targets the required elbow motion (see also Ahmadi-Pajouh et al, 2012). Given the re-organization of motor circuits during motor learning (Della-Maggiore et al, 2015; Doyon and Benali, 2005), it seems reasonable to suggest that long-latency stretch responses co-vary with motor learning because of the overlap of neural circuits that generate rapid feedback responses and voluntary motor commands.

It is important to note that the extensive practice of the Mirror Mapping condition produced a series of unexpected results for trials in the Veridical Mapping condition. For instance, we observed that long-latency stretch responses evoked in the WF were larger for OUT trials compared to IN trials, despite the fact that participants generated appropriate wrist extension movements for OUT trials in this mapping condition (Figure 6). This rapid feedback response did not support the intended and executed movement, and brings into question the idea that the neural circuits that generate the long-latency stretch response and voluntary motor commands are the same and/or perform the same computations. We also observed that long-latency stretch responses evoked in the WF for OUT trials were indistinguishable between the Veridical and Mirror Mapping conditions. One possibility is that participants learned to appropriately complete the Mirror Mapping trials by releasing a preplanned wrist flexion movement (i.e., triggered response: Crago et al, 1976) after a mechanical perturbation (Ravichandran et al, 2013), and this learned effect carried over to the Veridical Mapping block. In this case, the perturbation would have had to selectively release one of two preplanned actions, because the WF’s long-latency response was modulated by the target’s location even though participants could not have planned the appropriate response prior to the perturbation. With extensive training it may be possible to selectively release one of a several preplanned actions, as there is evidence that individuals can preplan two simultaneous movements within the trials of a single experimental session (Carlsen et al, 2009, but see Carlsen et al, 2004). In addition to two preplanned movements, there would also need to be another signal to override the preplanned wrist flexion response to rapidly attenuate the initial long-latency stretch responses observed in the WF (see Figure 7B) and allow participants to produce the appropriate wrist extension movement that was observed for trials in the Veridical Mapping condition. This is theoretically possible, as it has been proposed that the release of preplanned movements depend on brainstem circuits (Carlsen et al, 2004; Honeycutt and Perreault, 2012) whose output could be overridden by the descending cortical commands that generate voluntary movement.

